# PFAS exposure is associated with accelerated epigenetic ageing in a wild marine mammal

**DOI:** 10.64898/2026.05.29.728902

**Authors:** Katharina J. Peters, Karen A. Stockin, Eva-Maria F. Hanninger, Livia Gerber

## Abstract

Chronic contaminant exposure may impose hidden physiological costs long before obvious demographic or health effects become detectable in wildlife populations. Epigenetic clocks quantify biological ageing and may provide sensitive biomarkers of cumulative toxicological stress. Per-and polyfluoroalkyl substances (PFAS) are persistent contaminants that bioaccumulate in marine food webs, yet their long-term physiological consequences for wildlife remain poorly understood. Here, we tested whether PFAS exposure is associated with accelerated biological ageing in common dolphins (*Delphinus delphis*). We analysed liver PFAS concentrations and skin DNA methylation profiles from 30 stranded or bycaught dolphins from New Zealand waters. Epigenetic age was estimated using a recently developed species-specific epigenetic clock, and age acceleration was calculated as the residual deviation between epigenetic and chronological age. Using an information-theoretic modelling framework, we assessed the effects of total PFAS burden, sex, and their interactions on epigenetic age acceleration. Total PFAS concentrations were positively associated with epigenetic age acceleration, indicating that dolphins with higher PFAS burdens were biologically older than expected for their chronological age.

Each 1 ng g□¹ increase in total PFAS was associated with an average increase of 0.031 years in biological age. Sex did not significantly influence age acceleration, suggesting that PFAS-associated ageing effects occur across both sexes. Although modest, this effect is consistent with PFAS acting as a chronic physiological stressor influencing molecular ageing processes. Our findings provide the first evidence linking PFAS exposure to accelerated biological ageing in a wild mammal, highlighting epigenetic ageing as an integrative biomarker of long-term contaminant effects in wildlife.

## Introduction

Per-and polyfluoroalkyl substances (PFAS) are a large and diverse group of synthetic chemicals that have been extensively manufactured and used worldwide due to their unique physicochemical properties, including thermal stability, resistance to degradation, and water-and oil-repellent characteristics (Abunada et al. 2020; Gaines 2023). These properties have led to their incorporation into a wide range of industrial processes and consumer products, such as non-stick cookware, firefighting foams, and water-resistant textiles (Glüge et al. 2020). Under the current OECD (Organisation for Economic Co-operation and Development) definition, PFAS comprise more than 14,000 documented chemical structures, including perfluoroalkyl acids (PFAAs) and a wide array of precursor compounds that can transform into persistent end products in the environment (Ahrens et al. 2009; Ahrens and Bundschuh 2014). PFAS are now recognised as contaminants of emerging concern because of their environmental persistence, long-range transport, and growing evidence of adverse effects on wildlife and human health (Johnson et al. 2021; Sonne et al. 2021; Green et al. 2024). Due to their resistance to degradation, the world’s oceans are considered a major sink for PFAS (Yamashita et al. 2004; Bashir et al. 2020), where these compounds are assimilated by marine organisms, bioaccumulate in protein-and phospholipid-rich tissues, and biomagnify through food webs, particularly in long-lived, higher-trophic-level species such as marine mammals (Kelly et al. 2009; Tomy et al. 2009; Fair and Houde 2018).

PFAS are among a growing class of environmental pollutants recognised as external drivers of biological ageing, a universal process marked by progressive decline in physiological function over time (Nussey et al. 2013). While chronological age simply reflects the number of years an individual has lived, individuals of the same species, and even of the same chronological age, often differ substantially in their rate of functional decline (Levine et al. 2018). This variation highlights the distinction between chronological versus biological age, the latter representing an individual’s physiological condition and overall health status. Biological age captures how “old” an organism is in functional terms and can diverge from chronological age due to genetic, environmental, and social influences (Rutledge et al. 2022). Recent advances in molecular biology, particularly the development of epigenetic clocks based on predictable age-related changes in DNA methylation (Horvath and Raj 2018; Barratclough et al. 2021; Peters et al. 2023; Barratclough et al. 2025), have enabled biological age to be quantified across a wide range of species. By identifying DNA sites that gain or lose methyl groups in a predictable manner over the course of a species’ lifespan, these tools have transformed ageing research by allowing investigators to examine how external factors, including social bonds, shape the ageing process itself, rather than lifespan alone (Gerber et al. 2025).

Although PFAS have been in use since the 1940s (Cheng et al. 2025), a limited but growing number of studies have so far investigated their effect on ageing, likely because measuring biological age has only been available on a larger scale in recent years (Dutta et al. 2023). In humans, PFAS exposure has been associated with accelerated biological aging in adults (Zhao et al. 2024; Khodasevich et al. 2025; Yan et al. 2025), as well as the opposite (Chaney and Wiley 2023), highlighting the need for further longitudinal research. In animals, studies on PFAS have focussed on laboratory or domesticated individuals (Peritore et al. 2023), rather than wildlife. While there is an understanding of the general negative effects of PFAS on animal health (Sonne et al. 2021; Parolini et al. 2022), no information exists on the relationship between PFAS and biological age in wild animals.

Here, we explore the effect of PFAS contamination on biological age in a marine predator, the common dolphin (*Delphinus delphis*). Specifically, using an epigenetic clock, we test if individuals with lower PFAS concentrations were biologically younger, i.e., have a younger epigenetic age than expected for their chronological age, when compared to individuals with higher PFAS concentrations. This is the first study to explore this relationship in a wild animal, providing important insights into the effect of forever chemicals on aging.

## Methods

### Sample collection

We analysed skin and liver tissue of 30 common dolphins ranging from 4 to 26 years of age (14 males, 16 females) stemming from a 30-year tissue biobank (BIOCET) curated from stranded or bycaught common dolphins. Individuals were a priori aged between 4 and 26 years of age (refer to ‘Chronological and epigenetic age estimation’). We only included individuals older than 4 years to ensure that measured PFAS burdens reflected each animal’s own cumulative environmental exposure, rather than maternally derived contaminants transferred during gestation and lactation, which can inflate PFAS concentrations in very young individuals (Stockin et al. 2025).

We extracted and purified DNA using a Quick-DNA Miniprep Plus Kit (Zymo) following the protocol for solid tissue samples, and a DNA Clean & Concentrator Kit (Zymo), respectively, as outlined in Hanninger et al. (2025).

### PFAS analysis

Hepatic tissues were analysed by AsureQuality Laboratories, Wellington, NZ, using validated in-house methods and quality control procedures, as detailed in Stockin et al. (2021). Briefly, frozen liver samples were thawed at 2–5 °C and then brought to room temperature. The samples were homogenised using a Waring blender, after which an ∼ 1 g aliquot was taken from the homogenate and an internal standard mixture was added. After adding acidified acetonitrile, the aliquot was further homogenised with ceramic homogeniser pellets using a Merris Minimix vibrational shaker for 8 min at 50 Hz. An aliquot of the resulting extract was cleaned up by dispersive solid-phase extraction (SPE) employing graphitised carbon, C18 sorbent, and zirconia-coated silica. Prior to LC–MS/MS analysis, extracts were prepared in a 50:50 methanol–water solvent system. Because suitable blank samples were unavailable for determination of the limit of detection (LOD), the limit of reporting (LOR) was defined as the lowest calibration standard, corresponding to 0.5 ng g□¹.

Quality control (QC) procedures followed the requirements of the US DoD/DoE QSM 5.1.1 (2018). Each analytical batch included an ongoing precision and recovery (OPR) sample and an OPR blank, with OPR samples fortified with analytical standards at 1–2 times the method LOR. A reagent blank (containing no matrix but processed with all extraction reagents and steps) was also included in each batch. In addition, one submitted sample per batch was randomly selected for fortification with target analytes at the OPR concentration both before and after extraction to evaluate matrix effects on analyte recovery. Each batch further included a randomly selected duplicate sample for quality assurance.

PFAS analysis followed validated in-house ESI-LC-MS/MS methods, measuring the following compounds: PFBA, PFHxA, PFOA, PFNA, PFDA, PFUnDA, PFDoDA, PFTrDA, PFTeDA, PFHpS, L-PFHxS, L-PFOS, mono-PFOS, PFDS, PFOSA, FPePA (5:3 FTA). For PFOS and PFHxS, linear and branched isomers were quantified separately, with branched isomers further distinguished as di-methyl and mono-methyl forms. Total branched PFOS (B.PFOS) and total branched PFHxS (B.PFHxS) were calculated as the sum of their respective branched isomers. For statistical analysis, we calculated total PFAS as the sum of all analysed compounds per sample.

### Chronological and epigenetic age estimation

We determined the focal animals’ chronological age by counting growth layer groups in the dentine of thin, decalcified, and stained sections of teeth. All dental ages were assessed by at least two experienced readers. Detailed methods on dental age estimation are described in Palmer et al. (2023).

We inferred epigenetic age of individual dolphins using our recently published common dolphin species-specific epigenetic clock (Hanninger et al. 2025). In brief, we calibrated the epigenetic clock using methylation proportions (β values) at 37,492 CpG sites, with data generated utilising the mammalian methylation array (Arneson et al. 2022). Here, we use epigenetic age estimates from the best-performing epigenetic clock in Hanninger et al. (2025) developed using an elastic net regression with Leave-One-Out Cross-Validation (LOOCV) on a curated dataset of common dolphins with high-confidence dental age estimates (N = 63, Median Absolute Error (MAE) = 1.72, Pearson R = 0.91, R^2^ = 0.83).

To quantify age acceleration, we calculated AgeAccel for each individual, a metric widely adopted in epigenetic clock research (Teschendorff and Horvath 2025). Also known as ‘extrinsic age acceleration’ (Horvath et al. 2016), this measure is derived by regressing epigenetic age against chronological age and extracting the residuals (Krieger et al. 2023), capturing the degree to which an individual’s epigenetic age deviates from what would be predicted by their chronological age alone (Horvath 2013).

## Statistical analyses

We considered linear mixed-effects and linear models to examine the relationship between PFAS concentration and age acceleration. Prior to fixed effects selection, we determined the optimal random effects structure by comparing models with and without sampling year as a random effect.

We employed an information-theoretic approach to select the fixed effects. We fitted a global model containing all biologically plausible fixed effects measured (total PFAS, chronological age, sex) and their interactions. We included chronological age not solely to infer possible interactions but also to control for chronological age while isolating the effect of sex and total PFAS on epigenetic age, as recommended by Krieger et al. (2023). We used the MuMin package’s ‘dredge’ function to generate all possible hierarchically valid model combinations. We ranked the models using Akaike Information Criterion corrected for small sample sizes (AICc) (Hurvich and Tsai 1989), with lower values indicating better model fit. In our results, we present the results from the model with the lowest AIC. We verified the model assumptions through examination of residual plots, Q-Q plots of residuals, and Q-Q plots of random effects. Statistical significance was assessed at α = 0.05. All analyses were done using R version 4.5.2 (R Core Team 2025).

## Results

### Association between PFAS exposure and DNA methylation age acceleration

Our comparison of models with and without sampling year as a random intercept did not support the inclusion of this random effect (ΔAICc < 2), and thus, we did not include sampling year in subsequent analyses. Model selection using AICc identified the model including sex and total PFAS concentrations as predictors of age acceleration, without interaction terms or chronological age (Table 1).

**Table 1.** Model selection results (top five candidate models) for age acceleration in common dolphins (*Delphinus delphis*) in New Zealand. Candidate models were generated as all subsets of a global linear model (AgeAccel ∼ total PFAS * age * sex) using MuMIn::dredge and ranked by Akaike’s Information Criterion (AICc). ΔAICc is calculated relative to the lowest-AICc model, and Akaike weights (AICc_w_) represent the relative support for each model within the candidate set. *df* is the number of estimated parameters, including the intercept and residual variance (σ²). Total PFAS is expressed as ng/g wet weight.

| Model terms | df | AICc | $\Delta\text{AICc}$ | $\text{AICc}_w$ |
| --- | --- | --- | --- | --- |
| sex + total<br>PFAS | 4 | 135.7 | 0.00 | 0.209 |
| total pfas | 3 | 135.9 | 0.21 | 0.188 |
| null model | 2 | 136.2 | 0.51 | 0.162 |
| age | 3 | 137.3 | 1.62 | 0.093 |
| sex | 3 | 137.9 | 2.21 | 0.069 |

This best-supported model explained 17.2% of the variance in age acceleration (R^2^ = 0.172, p = 0.078). Total PFAS concentrations showed a significant positive association with age acceleration (β = 0.031, SE = 0.014, p = 0.038), indicating that dolphins with higher PFAS burdens exhibited accelerated ageing relative to their chronological age (Table 2, Fig 1). Specifically, each 1 ng/g increase in total PFAS was associated with a 0.031-year (approximately 11 days) increase in age acceleration. While sex did not significantly influence age acceleration (β =-1.354, SE = 0.820, p = 0.110, Table 2, Fig 1), it was included as a variable in the top-ranked model.

**Fig 1.**
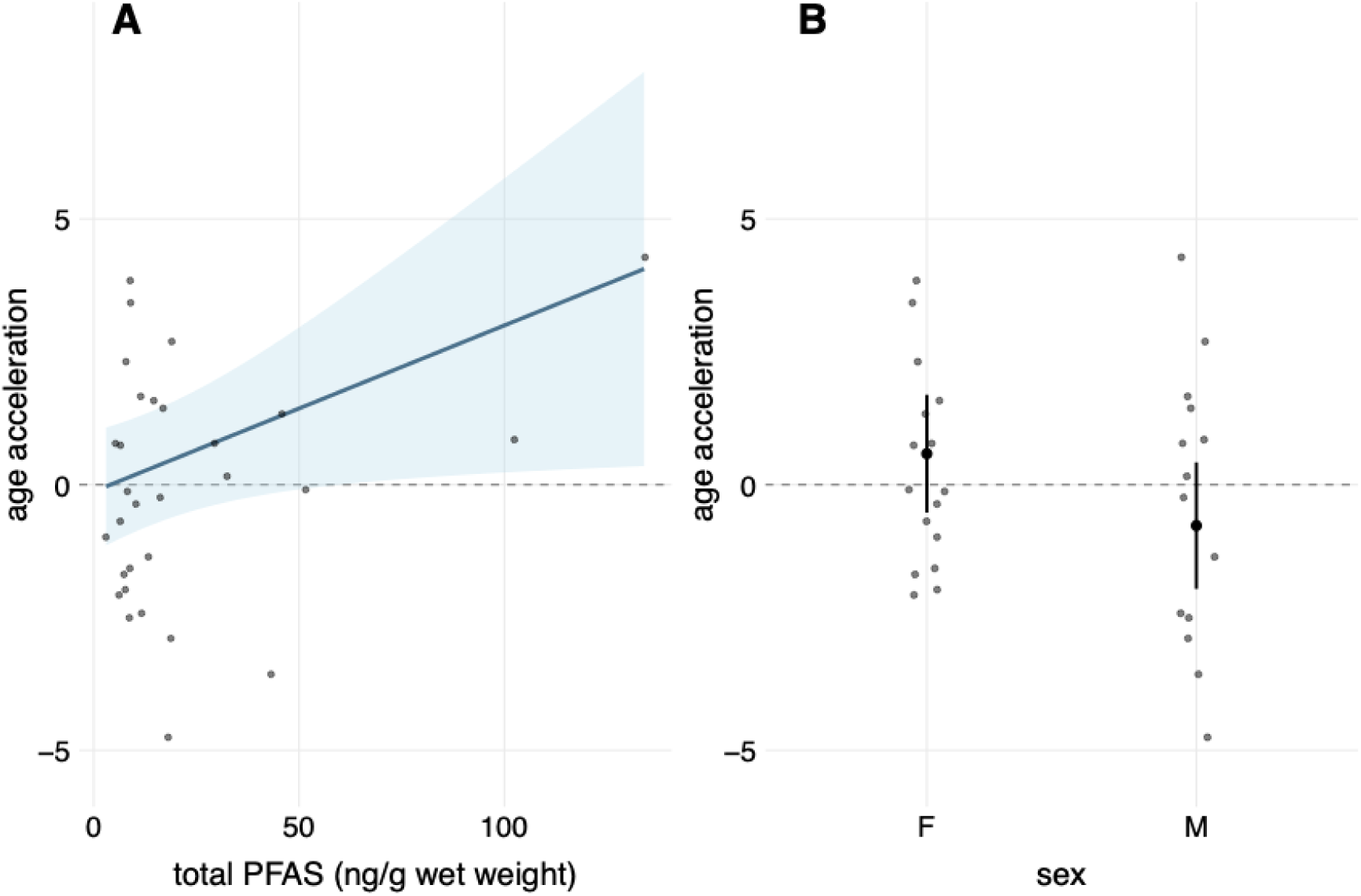
(A) Relationship between total PFAS concentrations (ng/g wet weight) and age acceleration in common dolphins (*Delphinus delphis*) in New Zealand. The blue line represents the fitted relationship from the best-supported linear model, with the shaded area indicating 95% confidence intervals. Points show individual dolphin observations (n = 30). Each 1 ng/g increase in total PFAS concentrations was associated with a significant 0.031-year increase in age acceleration (*p* = 0.038). (B) Effect of sex on age acceleration. Males showed reduced age acceleration compared to females, though this difference was not statistically significant (β = - 1.354, *p* = 0.110). Box plots show median, interquartile range, and individual data points. Age acceleration represents the deviation of epigenetic age from chronological age, with positive values indicating accelerated aging and negative values indicating decelerated aging.

**Table 2.**
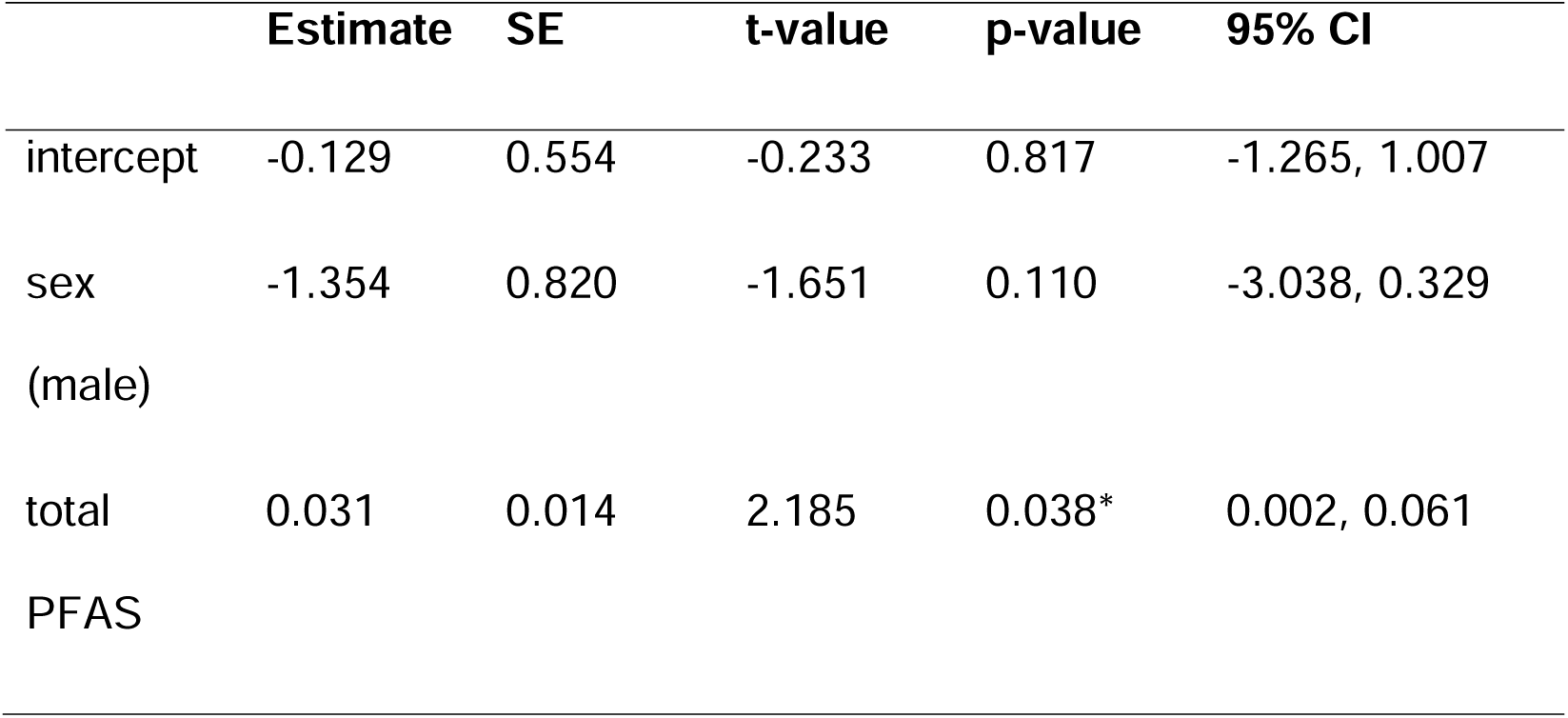
Best-performing linear model explaining age acceleration (AgeAccel) in common dolphins (*Delphinus delphis*) in New Zealand as a function of sex and total PFAS concentration (ng/g wet weight). Coefficients are unstandardized estimates (β) ± SE with p-values from t-tests. Model summary: N = 30; R² = 0.172; adjusted R² = 0.111; overall F(2,27) = 2.81, p = 0.078.

## Discussion

Here, we provide first insights to the effect of PFAS on biological age in a wild mammal. We found a positive association between total PFAS concentration and epigenetic age acceleration in common dolphins, meaning that individuals with higher total PFAS burdens exhibited greater biological age relative to their chronological age. This finding provides first evidence that PFAS exposure is associated with subtle but measurable changes in molecular ageing processes in a free-ranging marine mammal. While the magnitude of the effect was modest, epigenetic age acceleration reflects cumulative physiological stress and altered regulatory processes rather than acute toxicological responses, making it a sensitive indicator of long-term biological impact (Suglia et al. 2024).

PFAS compounds are known to disrupt endocrine signalling (Di Nisio et al. 2022; Bulawska et al. 2025), induce oxidative stress (Iakovou and Kourti 2022; Solan et al. 2023), and alter immune and metabolic pathways (Guo et al. 2022), all of which are processes closely linked to epigenetic regulation and ageing. Experimental and observational studies in other taxa have shown that chronic PFAS exposure can influence DNA methylation patterns (in humans; Schmidt 2022; Abdulkadir et al. 2025), mitochondrial function (in humans; Hofmann et al. 2023; Kam et al. 2025), and inflammatory responses (in humans and animals; Zhang et al. 2023; Bline et al. 2024), providing plausible mechanistic pathways through which PFAS may contribute to accelerated biological ageing. Although we did not assess the specific molecular mechanisms here, the observed association is consistent with PFAS acting as a chronic physiological stressor capable of influencing epigenetic ageing trajectories.

The observed effect size of PFAS on epigenetic age acceleration was relatively small, with increases in PFAS concentration corresponding to incremental increases in biological age. This is not unexpected given that epigenetic ageing integrates multiple intrinsic and extrinsic influences accumulated over an individual’s lifetime (Pal and Tyler 2016; Navarro et al. 2023; Zhang et al. 2023). Ageing processes are shaped by a complex interplay of genetics, environmental conditions, nutritional status, disease exposure, and contaminant load, and PFAS exposure represents only one component of this broader context. Consequently, even modest shifts in epigenetic age may reflect meaningful biological costs when sustained over long timeframes.

The modest magnitude of the association may also indicate that common dolphins possess physiological mechanisms that partially buffer or mitigate the effects of PFAS exposure over their lifespan. The liver plays a central role in PFAS uptake, sequestration, redistribution, and elimination kinetics through protein binding and enterohepatic circulation rather than classical biotransformation (Lu et al. 2024; Fischer et al. 2025), and individuals may differ in their capacity to sequester and redistribute PFAS compounds over their lifespan. Such variation could reduce the detectable impact of PFAS on ageing at the population level while still imposing costs on more susceptible individuals. Additionally, long-lived marine mammals may experience gradual PFAS elimination or redistribution among tissues (Sciancalepore et al. 2021; Khan et al. 2023), potentially dampening age-related accumulation and its downstream effects. It is also worth noting that the epigenetic clock used here was calibrated on skin tissue. A clock calibrated on hepatic tissue may have been better suited to detect associations between PFAS and epigenetic ageing, given the liver’s central role in PFAS uptake and sequestration. Skin, as a peripheral tissue exposed to external environmental pressures such as salinity and UV radiation, may introduce additional variation into methylation estimates that is unrelated to internal contaminant burden. Nevertheless, peripheral tissues have been shown to reflect biologically meaningful epigenetic signals associated with physiology and environmental exposure (Crossman et al. 2021; Mancia et al. 2021), suggesting that skin-based clocks remain a valid, if imperfect, tool for investigating contaminant-ageing relationships in wild cetaceans where tissue access is inherently constrained.

Despite potential buffering mechanisms, the association between PFAS burden and epigenetic age acceleration suggests that PFAS exposure may contribute to cumulative physiological wear over time. In long-lived species such as dolphins, even small annual increases in biological ageing could translate into reduced health, altered reproductive performance, or decreased resilience to additional stressors such as climate-driven prey shifts, disease outbreaks, or other contaminants.

Importantly, the broad occurrence of PFAS-associated ageing across individuals indicates a population-wide exposure risk rather than an effect confined to a particular demographic group.

In addition to PFAS exposure, we evaluated whether epigenetic age acceleration differed between sexes. Although sex was retained in the best-supported model, its effect on epigenetic age acceleration was not statistically significant, and the associated confidence interval overlapped zero. The negative coefficient suggests that males tended to exhibit lower age acceleration than females, but the uncertainty around this estimate indicates that any sex-specific difference in epigenetic ageing is weak relative to the effect of PFAS burden. Sex differences in contaminant toxicokinetics, reproductive investment, and hormone regulation could plausibly influence ageing processes in marine mammals.

In odontocetes, females are known to experience contaminant offloading during gestation and lactation (Lee et al. 2022), while males may accumulate contaminants more steadily over time. Previous studies have found varying sex-related differences in PFAS burden, with no difference in PFAS levels between male and female white-sided dolphins (*Lagenorhynchus acutus*) in the St. Lawrence Estuary, Canada (Khalid et al. 2024), but higher concentrations in males across a range of odontocete species (Stockin et al. 2025; Stokes et al. 2026), which matches the female-offloading dynamic. Despite these differences, our results suggest that PFAS-associated epigenetic ageing occurs broadly across both sexes, reinforcing the interpretation that contaminant exposure represents a population-wide physiological stressor rather than one restricted to a particular demographic group.

The best-supported model explained 17.2% of the variance in epigenetic age acceleration, consistent with ageing being a multifactorial process influenced by intrinsic and extrinsic factors not captured in this study. This study is based on cross-sectional data, which limits causal inference and prevents assessment of individual ageing trajectories through time. Future work incorporating longitudinal sampling, compound-specific PFAS profiles, and additional physiological markers such as sexual maturity status, would help to clarify the mechanisms linking PFAS exposure to epigenetic ageing and identify thresholds beyond which biological impacts become more pronounced (Stockin et al. 2025). Integrating epigenetic ageing with measures of health, reproduction, and survival will be particularly important for assessing the long-term population-level consequences of PFAS exposure in marine mammals. Nevertheless, this study provides novel, ecologically relevant evidence that PFAS exposure is associated with accelerated biological ageing in a wild marine mammal. More broadly, our findings highlight epigenetic age acceleration as a promising integrative biomarker for detecting the hidden physiological costs of chronic contaminant exposure in free-ranging wildlife.

## Acknowledgements

This research was supported by a University of Wollongong Environmental Futures Enable Grant to KJP, LG and KAS, and Massey University Research Funding (KAS). KAS was supported by a Royal Society Te Aparangi Rutherford Discovery Fellowship.

## Data availability statement

Data will be made available on GitHub.

## Author contributions

Katharina J. Peters: Conceptualization, Methodology, Formal analysis, Funding acquisition, Investigation, Data curation, Visualization, Writing – original draft, Writing – review & editing.

Livia Gerber: Conceptualization, Methodology, Formal analysis, Funding acquisition, Investigation, Visualization, Writing – original draft, Writing – review & editing.

Karen A. Stockin: Conceptualization, Resources, Data curation, Laboratory analyses, Funding acquisition, Sample collection, Writing – review & editing.

Eva Maria F. Hanninger.: Laboratory analyses, Data curation, Writing – review & editing.

